# Sex-dependent neuronal loss and apoptosis-associated signaling in the anterior cingulate and anterior insular cortices in a late-stage MIA mouse model of osteoarthritis

**DOI:** 10.64898/2026.02.09.704883

**Authors:** Oumaima Moutayb, Jaques Noel, Youssef Anouar, Mohammed Bennis, Saadia Ba-m’Hamed, Lamghari Moubarrad Fatima-Zahra

## Abstract

Osteoarthritis (OA) is a leading cause of disability worldwide, with chronic pain representing its most debilitating symptom and frequently accompanied by affective and cognitive comorbidities. Increasing evidence implicates maladaptive supraspinal plasticity within cortical regions involved in pain affect, including the anterior cingulate cortex (ACC) and anterior insular cortex (AIC), however, the relationship between these behavioral impairments and neuronal alterations, as well as potential sex-specific vulnerability, remains poorly documented. Using a monosodium iodoacetate (MIA) model of knee OA in adult male and female mice, we examined the temporal progression of sensory, affective, and cognitive alterations at early (day 7) and advanced (day 28) stages of disease. Pain sensitivity, locomotor and gait changes, anxiety- and depression-like behaviors, and working-memory performance were assessed using established behavioral paradigms, followed by analyses of apoptosis-associated neuronal signaling in the ACC and AIC. MIA induced robust mechanical and thermal hypersensitivity and gait impairment in both sexes, while early emotional and cognitive alterations were not observed. In contrast, advanced OA was associated with pronounced anxiety- and depression-like behaviors and impaired working memory. Notably, analysis at day 28 post-MIA revealed a significant increase in apoptotic signaling and neuronal loss in both cortical regions, with females exhibiting greater vulnerability, particularly within the AIC, paralleling their more severe affective phenotypes. Together, these findings indicate that chronic OA pain is associated with progressive, sex-dependent neuronal loss within key cortical pain-affective circuits and highlight supraspinal remodeling as a potential substrate underlying the emotional and cognitive burden of OA pain.

## Introduction

The rapid global increase in life expectancy is shifting demographics toward older populations, with individuals aged 65 and above projected to represent nearly one-sixth of the world’s population by 2050 (WHO, 2024). This transition underscores the urgent need to address age-related health conditions that significantly impair quality of life. Among these, osteoarthritis (OA), a major musculoskeletal disorder, stands out as the most prevalent degenerative joint disease, affecting more than 50% of the elderly population worldwide (WHO, 2023). Beyond its clinical burden, OA imposes substantial socioeconomic consequences due to loss of independence, reduced productivity, and escalating healthcare costs (Hiligsmann *et al*. 2013).

Traditionally viewed as a purely mechanical “wear-and-tear” disease restricted to cartilage breakdown, OA is now recognized as a multifactorial whole-joint pathology driven by persistent low-grade inflammation, nociceptor sensitization, and neuromodulatory alterations within the central nervous system (Loeser *et al*. 2012; McDougall, 2019). Chronic pain is the most disabling symptom of OA, representing the foremost reason for clinical intervention (Dieppe and Lohmander, 2005). Importantly, its severity does not correlate with radiographic joint damage, strongly implicating central mechanisms in pain persistence (Suokas *et al*. 2012).

Over time, OA pain evolves from an intermittent, nociceptive state into a chronic, self-sustaining condition characterized by central sensitization, maladaptive plasticity, and dysregulation of pain–emotion circuitry (Thakur *et al*. 2014; Meints and Edwards, 2018). Consistent with this, patients frequently exhibit cognitive impairments and affective disturbances (including anxiety, depression, and memory deficits) that exacerbate disease burden and lead to a poorer prognosis (Blackburn-Munro and Blackburn-Munro, 2001; Zhuo, 2016). These features are mediated by aberrant function of cortical regions integrating sensory, emotional, and interoceptive dimensions of pain, especially the anterior cingulate cortex (ACC) and anterior insular cortex (AIC) (Russell *et al*. 2018; Pujol *et al*. 2022).

Neuroimaging studies support a direct involvement of these regions in OA neurobiology, reporting reduced gray-matter volume, altered cortical thickness, and dysfunctional connectivity within ACC and AIC networks (Salazar-Méndez *et al*. 2023). Moreover, these structural and functional changes often correlate with symptom severity and may predict poor surgical outcomes (Soni *et al*. 2019). However, whether these cortical abnormalities reflect reversible plastic adaptations or irreversible neuronal loss remains largely unresolved, particularly at the cellular level.

Sex-specific differences further complicate OA pathophysiology. Women exhibit a markedly higher OA incidence, faster progression, and more severe pain and comorbid anxiety/depression than men (Glass *et al*. 2014; Sharma *et al*. 2016; Segal *et al*. 2024). Despite this growing evidence, the neuronal basis and behavioral expression of sex differences in OA remain poorly characterized. Many preclinical studies have historically focused on male animals, leading to a substantial knowledge gap regarding female vulnerability and the temporal progression of pain and comorbid symptoms.

To address this critical gap, we used the MIA-induced OA model in both male and female mice to longitudinally assess spontaneous, affective, and cognitive disturbances across disease progression. We then tested whether these behavioral impairments are associated with caspase-3–mediated apoptosis and neuronal loss in the ACC and AIC.

This integrated behavioral and histological approach aims to reveal sex-specific vulnerability to supraspinal neuronal alterations during OA, thereby refining mechanistic understanding and guiding the development of more personalized therapeutic strategies.

## Materials and Methods

### 1. Animals

In this study, young adult (3 months old; 25-30 g) male and female *Swiss* strain mice (Cadi Ayyad University, Marrakech, Morocco) were used, housed in Plexiglas cages (30 cm × 15 cm × 12 cm), under standard conditions (22 ± 2 °C; 12 h/12 h light/dark cycle), with *ad libitum* access to food and water except during behavioral testing. All procedures were performed in accordance with approved institutional protocols and the provisions for the care and use of animals in scientific procedures set out in the European Council Directive EU2010/63. Every effort was made to minimize animal suffering. The study was approved by the Council Committee of Research Laboratories of the Faculty of Science, Cadi Ayyad University, Marrakech (BA-05/2024).

### 2. Induction of knee osteoarthritis

Mice were randomized in two groups, control mice (sham, saline-treated; male, n = 27; female, n = 27) and osteoarthritis mice (OA, MIA-treated; male, n =27; female, n =27).

At day 0 corresponding to postnatal day (PND) 90, mice were anesthetized with an intraperitoneal (IP) injection of ketamine (5 mg/kg)/xylazine (2 mg/kg), 1 mL/kg body weight. They then received a single IA injection into the right knee of 1 mg MIA (Sigma-Aldrich) in a total volume of 10 μL of physiological sterile saline using a microsyringe (26 G) (Pitcher *et al*. 2016). The limbs of the mice were delicately secured to an operating table with hemp ropes while lying flat. Their knee joints were flexed at 90° to expose the patellar ligaments, and needles were inserted through the parapatellar joint space. MIA injections require great care to avoid leakage of the substance from the articular capsule, which would cause systemic toxicity resulting in animal death. Mice in the control group received an IA injection of an equivalent volume of saline.

### 3. Behavioral testing and study design

Mice were acclimated to the testing environment for 30 minutes prior to behavioral assessments. Baseline behaviors were recorded on day 0 (PND90), before the OA induction. Two experimental series were conducted to evaluate both the OA phenotype and its associated comorbidities. Assessments of inflammation and pain-related behaviors were conducted at two time points: 7 days post-MIA injection (PND97) to represent the early stage of OA progression, and 28 days post-MIA injection (PND118) to represent the late stage Figure 1). To investigate OA-related comorbidities, including anxiety-like behavior, depressive-like behavior, and working memory, behavioral tests were also administered on days 7, and 28 post-injections. To minimize potential habituation effects during the second experiment, different groups of animals were used at each time point. All behavioral evaluations were performed in the morning (between 9:00 AM and 12:00 PM) under standardized lighting conditions. Behavioral activity was recorded using EthoVision XT software (Noldus, The Netherlands). All apparatuses were cleaned with 70% ethanol between trials, to eliminate confounding influences of residual odors. Hormonal status of female mice was not controlled. However, this approach is consistent with current recommendations in behavioral neuroscience, as synchronizing the estrous cycle is not required for valid comparisons and can even introduce artificial hormonal states. Several studies (Becker *et al*. 2005; Prendergast *et al*. 2014) demonstrate that natural estrous cycling does not increase variability compared with males and does not invalidate behavioral or neurobiological findings.

**Figure 1:**
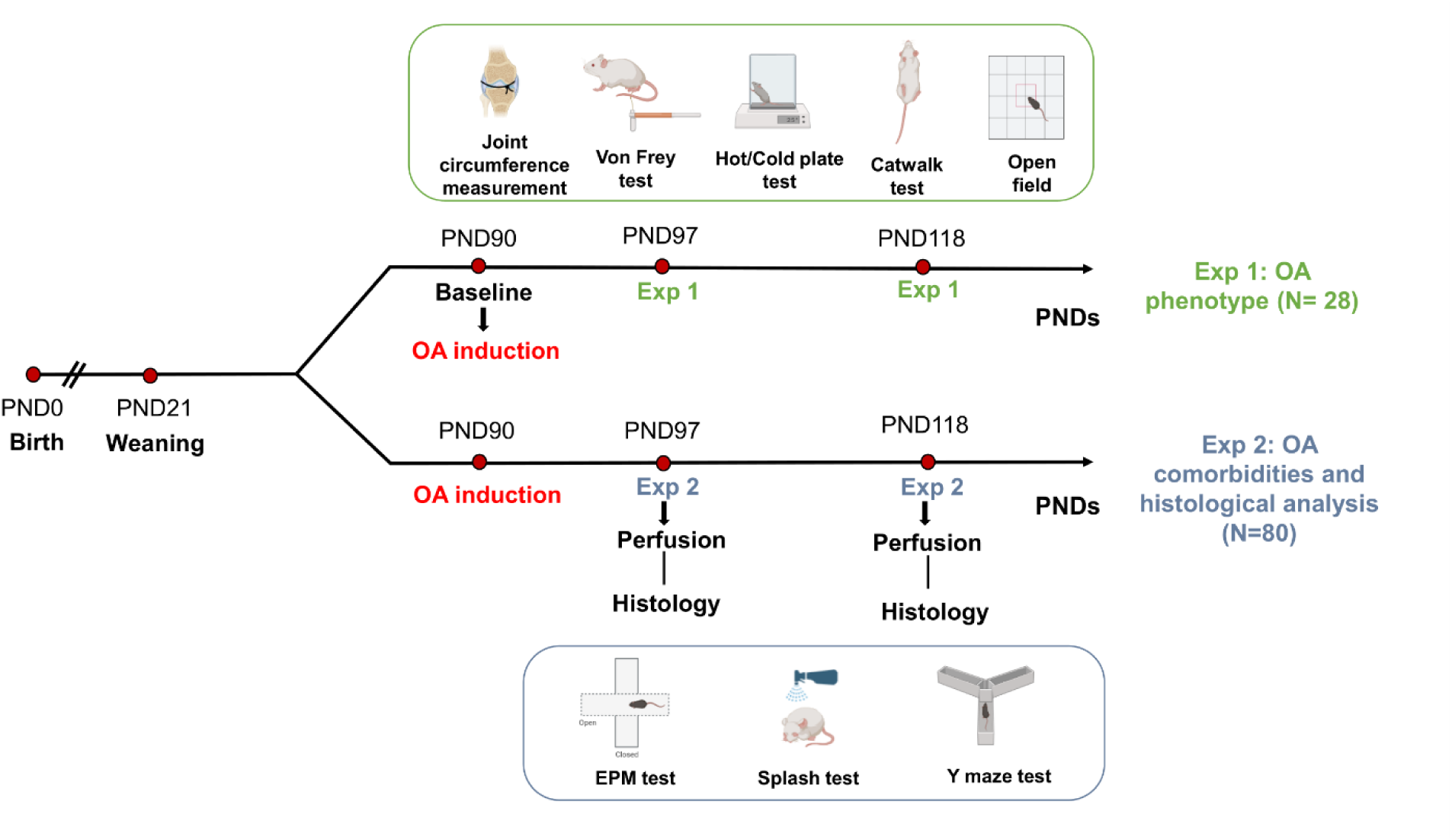
Experimental design of the study.

Following the behavioral assessments, histological analyses were performed to examine neuronal changes at 7- and 28-days post-injection. Neuronal density was quantified on Nissl-stained sections, and the apoptotic index was assessed by double immunofluorescence for NeuN and c-Caspase-3 in the contralateral (left) ACC and AIC (Figure 1).

#### 3.1. Joint swelling assessment

To evaluate the inflammatory response following the IA injection of MIA, we quantified the extent of knee swelling. By measuring the circumference of the joint using a thread. The mice were initially restrained gently to gain access to the knee. Subsequently, a non-stretchable thread was wrapped around the widest part of the ipsilateral knee joint, taking care to avoid indenting the skin. The point of intersection between the thread and the skin was marked, and the length up to the mark was measured using a ruler to obtain the circumference in centimeters. This procedure was repeated for both groups at designated time points post-injection to evaluate changes over time.

#### 3.2. Mechanical and thermal pain

##### i. Von Frey test

To assess mechanical allodynia, mice were individually housed in a plastic cage (12 × 12 × 10 cm) with a mesh floor to allow access to the paw. After a 1 h acclimation, mechanical sensitivity was assessed with calibrated von Frey filaments (0.02–8.0 g). Each filament was applied seven consecutive applications to the plantar surface of the hind paw ipsilateral to the IA-injected knee in both MIA- and saline-treated mice, using enough force to induce filament bending. This was then maintained for 3 s or until the paw was withdrawn in a movement not associated with a reflex or grooming, like flinching, paw lifting, or shaking (Chaplan *et al*. 1994).

The withdrawal threshold was determined as the filament at which the mice withdrew the paw at least four times in seven applications. Paw withdrawal thresholds (PWT) were expressed in grams (g).

##### ii. Hot / Cold plate test

To evaluate thermal sensitivity, hot and cold plate tests were conducted as previously described (Tjølsen *et al*. 1991; Jasmin *et al*. 1998). Mice were individually placed in a 20 cm-high transparent Plexiglas cylinder positioned on a metal surface maintained at either 55 ± 1 °C (hot plate) or 5 ± 1 °C (cold plate). To avoid tissue damage, animals were immediately removed from the testing surface upon the first observable nociceptive response or after a maximum exposure (cut-off) time of 30 seconds. Nociceptive behavior was defined as a rapid withdrawal or licking of the ipsilateral hind paw, and these responses were recorded as indicators of thermal threshold.

#### 3.3. Functional / spontaneous pain - related behaviors

##### i. Catwalk test

A 40 cm long, 5 cm wide runway and bounded by walls of 15 cm, was lined with white paper at the bottom. Mice were trained to run down the runway in a straight line a day before the test. On the test day, the paws of the mice were dipped in non-toxic acrylic paint and mice were allowed to run down the runway. The measured parameter is stride length of ipsilateral hind paw (distance covered in one complete walking cycle by the same hind paw) as an indicator of pain - related gait alteration and discomfort (Lakes and Allen, 2016).

##### ii. Open field test (OF)

To evaluate the changes of locomotor and exploratory activity in OA mice, OF test was performed as previously described (Wilson *et al*. 1976). It consisted of a square Plexiglas box (50 × 50 × 50 cm) in which the animal was placed in the center zone of the apparatus. Animal behavior was recorded for 10 min, during which, the total distance moved, animal mean velocity and the time spent in the center were assessed.

#### 3.4. Anxiety-like behavior

##### i. Elevated plus maze (EPM)

This assessment constitutes a validated method for examining anxiety-like behavior in rodents (Handley and Mithani, 1993). The apparatus comprises a cross-shaped maze constructed from transparent Plexiglas, featuring two open arms measuring 50 × 5 cm each, and two enclosed arms (50 × 5 × 15 cm), connected by a central platform (5 × 5 cm). The apparatus is positioned 45 cm above the floor. To initiate each trial, mice were introduced to the central area, facing one of the open arms, and permitted unrestricted exploration for a period of 5 min. The anxiety index (AI = 1 − [(open arm time/total time) + (open arm entries/total entries)] / 2) was recorded for each animal (Rao and Sadananda 2016).

#### 3.5. Depression-like behavior

##### i. Splash test

The aim of the splash test is to gain insight into depression-like symptoms in mice, such as self-care issues and motivational impairments (David *et al*. 2009). The test involves gently spraying a 10% sucrose solution onto the dorsal coat of each mouse in its home cage. Following the application of the sucrose solution, the time spent grooming was recorded for a period of five minutes.

#### 3.6. Working memory assessment

##### ii. Y-maze test

The experiment was conducted in a Y-shaped maze consisting of 3 identical arms, each measuring 10 cm in width, 41 cm in length, and 25 cm in height, spaced 120 degrees apart. Each mouse was subjected to a single trial, during which it was placed individually at the end of one of the maze’s arms and permitted eight minutes for exploration (Hughes, 2004). A successful spontaneous alternation was defined as the mouse entering all three arms consecutively without repeating any arm. The percentage of spontaneous alternation for each mouse was calculated using the following formula: % of spontaneous alternation = 100 x [(number of alternations) / (total arm entries - 2)].

### 4. Ex vivo assays

Following behavioral tests, animals were randomly assigned to Nissl staining or immunohistochemistry experiment.

#### 4.1. Nissl staining of brain slices

Once the behavioral testing had been completed, we performed a histological Nissl staining on brain slices. Mice were anesthetized with an intraperitoneal injection of ketamine/xylazine (5–2 mg/kg, 1 mL/kg body weight) and transcardially perfused with saline solution (0.9%), followed by 4% paraformaldehyde (PFA) in phosphate buffered saline (PBS; 0.1 M). The brains were gently removed from the skull, post-fixed in the same fixative for 12 h. Next, the contralateral hemispheres (left) were sectioned into 40 μm frontal sections using a vibratome (Leica VT1200, Germany). The brain slices were mounted on gelatin-coated microscope slides and Nissl stained (0.1% Cresyl violet, 0.1% acetic acid, and 0.01 M sodium acetate in sterile water), followed by dehydration in a graded series of ethanol, clearing in xylene, and cover-slipping with Eukitt. Images from the 5 slides per animal contain ACC (chosen between AP 0.14 and AP 1.10) and AIC (chosen between AP 1.9 and AP 2.2) were acquired using an Olympus BH-2 microscope, which is equipped with a camera (Olympus DP71). The methodology for counting cell density was based on the approach previously described by Van Heukelum *et al*. (2021).

#### 4.2. Immunohistochemistry experiments

In order to confirm this neuronal loss and to detect apoptotic neurons, a double immunofluorescence was conducted. Mice were anesthetized with an intraperitoneal injection of ketamine/xylazine (5–2 mg/kg, 1 mL/kg body weight) then transcardially perfused. Contralateral hemispheres were collected. Double immunostaining of NeuN (neurons) and c-Caspase-3 (apoptosis marker) was done on 40 μm thick brain floating sections containing the regions of interest (5 slides/ animal), according to stereotaxic atlas of Franklin and Paxinos (2001). Anti-mouse NeuN (1:100, #MAB377, Sigma-Aldrich), Anti-Rabbit c-Caspase-3 (1:800, 236 003, SYSY) were used as primary antibodies. Alexa 488 goat anti-mouse and Alexa 594 goat anti-rabbit were used as secondary antibody (1:1000, #A-11008 and #A-11012, Invitrogen). Sections were mounted in Vectashield solution (H-1000, Vector Laboratories). Images mosaics for cell counting were acquired with an inverted epifluorescence microscope (Axiovert 200 M, Carl Zeiss, Rueil Malmaison, France) through a 10x/0,3 objective. Cell counting was done with ImageJ / Fiji software (Schneider *et al*. 2012) on image of ACC and AIC. Three parameters were extracted: density of c-Caspase-3^-^/NeuN^+^ neurons, density of c-Caspase-3^+^/NeuN^+^ neurons and an apoptotic index was calculated as the proportion of NeuN⁺ neurons co-expressing c-Caspase3: AI=(cCasp3^+^/NeuN^+^)/(cCasp3^-^/NeuN^+^ + cCasp3^+^/NeuN^+^)×100% (Bressenot *et al*. 2009). To illustrate the apoptotic neurons, images from ACC of a female MIA were acquired with a Laser Scanning Confocal Microscope (TCS SP8, Leica, Microsystems, France) through a 63x/1.4 oil immersion objective with a voxel size of 0.09 x 0.09 x 0.20 μm. 3D reconstruction was performed using Imaris® 9.6.1 software (Oxford Instruments, Belfast, UK).

### 5. Statistical analysis

Statistical analyses were performed using GraphPad Prism software (version 10.6.1). Data normality was assessed using the Shapiro–Wilk test. For OA phenotype parameters assessed longitudinally in the same animals, a three-way mixed ANOVA (treatment × sex × time) with repeated measures on the time factor was applied. In contrast, for OA comorbidities and histological analyses, where independent cohorts were used at each time point, a three-way ANOVA (treatment × sex × time) was performed with time treated as a between-subjects factor. *Post hoc* comparisons were conducted using the Holm–Sidak method to correct for multiple testing. All data are presented as mean ± standard error of the mean (SEM), and a *p*-value < 0.05 was considered statistically significant.

## Results

### 1. IA injection of MIA induces joint swelling in both sexes

Our results demonstrated that the IA injection of MIA induced significant swelling in the ipsilateral knee joint of both sexes of MIA-treated mice compared to the control group (Figure *2A*). The three-way ANOVA (treatment × sex × time) RM on time analyses on knee circumference following IA MIA injection revealed a significant effect of treatment and time [F_(1, 24)_ = 52.01, *p* < 0.001; F_(2, 48)_ = 137.7, *p* < 0.001, respectively]. The analysis also revealed a significant effect of the time × treatment [F_(2, 48)_ = 122.3, *p* < 0.001] and sex × treatment [F_(1, 24)_ = 9.55, *p* = 0.005] interactions. The Holm-Sidak *post hoc* analysis revealed that MIA IA injection produced a clear increase in knee circumference on day 7 in both male and females mice compared to their control groups (t = 8.03; t = 11.38, *p* < 0.001, respectively), This increase was greater in MIA-treated females than in MIA-treated males (t = 3.68, *p* = 0.022). By day 28, this treatment effect persisted in MIA-treated females compared to its control group (t = 5.5, *p* < 0.001) but was not evident in males (t = 2.2, *p* = 0.71). However, no sex difference was detected at this time point (t = 1.8, *p* = 0.81) (Figure *2A*). Our results highlight the importance of studying OA progression overtime and by sex.

**Figure 2:**
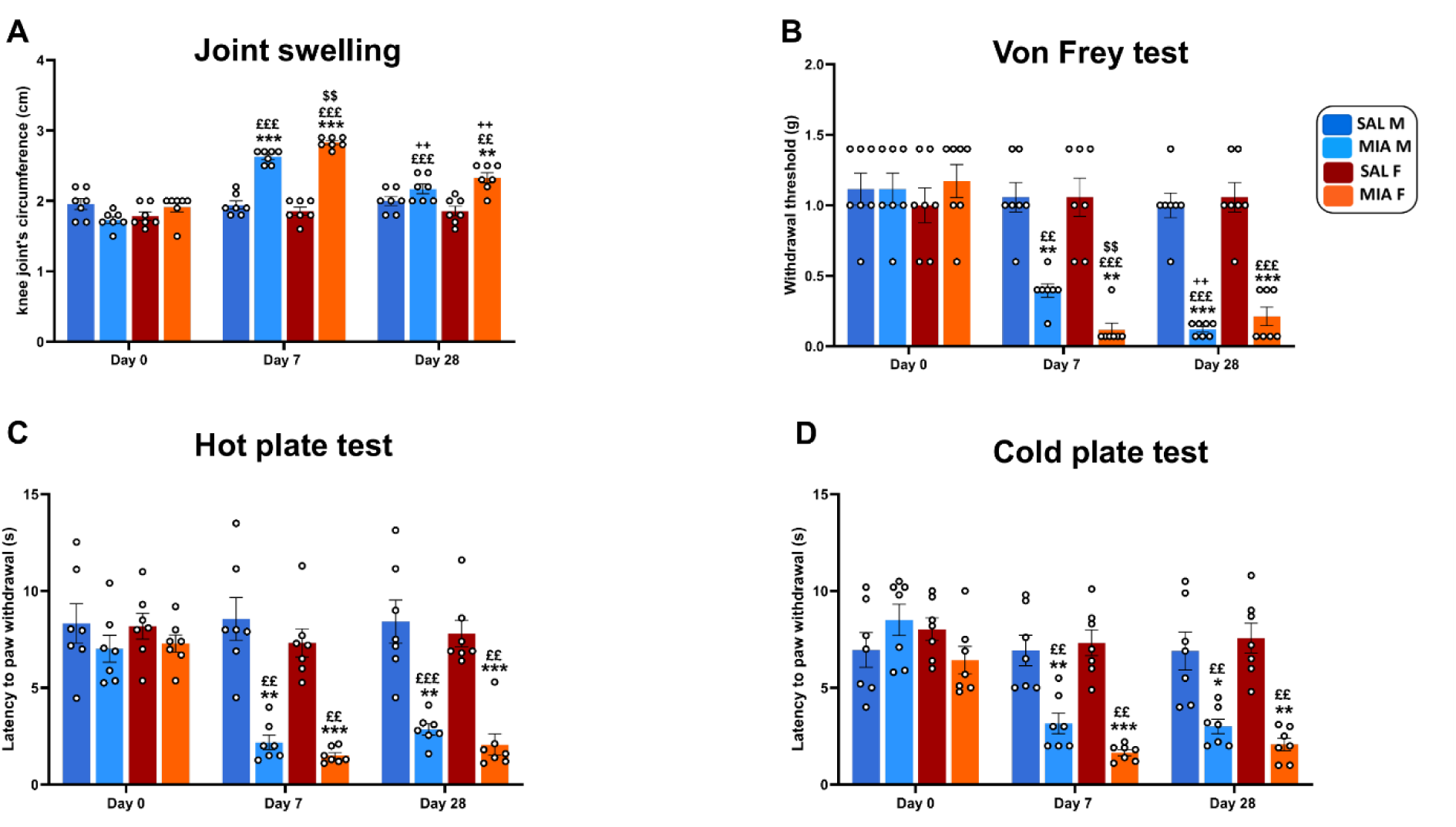
Time course of knee joint’s circumference, mechanical allodynia and thermal hypersensitivity development following IA MIA injection. **A:** Knee joint’s circumference. **B:** Paw withdrawal threshold using von Frey filaments. **C:** Paw withdrawal latency in the hotplate (55 °C). **D:** Paw withdrawal latency in the cold plate (5 °C). Results are represented as mean ± SEM (n=7/group). Holm-Sidak *post hoc*: \**p* < 0.05; ** *p* < 0.01; *** *p* < 0.001 *vs* same sex SAL; ££ *p* < 0.01; £££ *p* < 0.001 *vs* same sex MIA group at day 0; ++ *p* < 0.01 *vs* same sex MIA at day 7; $$ *p* < 0.01 sex difference.

### 2. IA injection of MIA induces mechanical allodynia in both sexes

Mechanical hypersensitivity is a key indicator of nociceptive dysfunction in OA. We therefore first examined whether IA MIA injection altered mechanical withdrawal thresholds across time in males and females by using Von Frey filaments. Three-way ANOVA (treatment × sex × time) RM on time revealed a significant effect of both treatment and time on withdrawal thresholds [F_(1,24)_ = 42.57; F_(2,48)_ = 69.50, *p* < 0.001, respectively], confirming a strong MIA-induced allodynia that evolved over the testing period. A significant interaction was also found between time and treatment [F_(2,48)_ = 64.82, *p* < 0.001] (Figure *2B*). Holm–Sidak’s *post hoc* multiple comparisons showed lower withdrawal thresholds in MIA-treated mice males and females on day7 compared to their control groups (t = 5.76, *p* = 0.0014; t = 6.52, *p* = 0.0013, respectively). Remarkably, MIA-treated females presented lower mechanical withdrawal compared to MIA-treated males (t = 4.11, *p* = 0.004), suggesting an early sex-dependent sensitivity to MIA. By day 28 post MIA injection both male and female mice still demonstrated lower withdrawal thresholds compared to control groups (t = 9.85; t = 6.83, *p* < 0.001, respectively), with no sex difference revealed (t =1.30, *p* = 0.41).Together, these findings show that IA MIA injection produces a strong and long-lasting mechanical allodynia, with females showing a more pronounced early response, but similar levels of hypersensitivity at later stages (Figure *2B*).

### 3. IA injection of MIA induces thermal hypersensitivity in both sexes

To further characterize thermal hypersensitivity alterations in both sexes of this model, we assessed sensitivity to heat and cold stimuli using the hot plate and cold plate tests. Both analyses were performed using a three-way RM ANOVA (treatment × sex × time) on time.

Analysis of the hot plate test (Figure *2C*) demonstrated significant effects of treatment, time and time × treatment interaction on latency to paw withdrawal [F_(1, 24)_ = 43.81; F_(2, 48)_ = 64.61 and F_(1, 24)_ = 53.31, *p* < 0.001, respectively]. However, neither sex, time× sex, sex × treatment nor time × sex × treatment interactions had significant effect [F_(1, 24)_ = 0.67, *p* = 0.41; F_(2, 48)_ = 1.97, *p* = 0.14; F_(1, 24)_ = 0.04, *p* = 0.83; F_(2, 48)_ = 0.25, *p* = 0.77, respectively]. The cold plate test (Figure *2D*) showed significant effects of treatment, time, and their interaction on latency to paw withdrawal [F_(1, 24)_ = 30.75; F_(2, 48)_ = 46.94; F_(2, 48)_ = 36.54, *p* < 0.001, respectively]. However, the factor sex, time× sex, sex × treatment and time × sex × treatment interactions had no significant effect on latency to paw withdrawal [F_(1, 24)_ = 0.48, *p* = 0.49; F_(2, 48)_ = 0.27, *p* = 0.76; F_(1, 24)_ = 3.80, *p* = 0.06; F_(2, 48)_ = 0.82, *p* = 0.44, respectively]. In both tests, Holm–Sidak’s *post hoc* revealed significant decrease of latency to paw withdrawal in MIA-treated male and female mice compared to control groups at both day 7 and day 28 with no sex differences. These findings demonstrate that IA MIA injection induces significant and sustained thermal hypersensitivity, to both heat and cold stimuli, independent of sex (Figure *2C* and D).

### 4. IA injection of MIA induces gait alteration and hypoactivity in both sexes

Because the persistence of pain often leads to measurable disruptions in motor function, we next investigated whether MIA-induced hypersensitivity was accompanied by changes in gait patterns and locomotor activity. Gait analysis and open-field testing were used to evaluate how pain-related discomfort affects spontaneous motor behavior over time in male and female mice.

Stride length, a sensitive indicator of gait alterations associated with joint pain, was markedly affected by MIA treatment (Figure *3A* and B). The three-factor, treatment × sex × time, ANOVA test RM on time showed significant effects of treatment, time, and their interaction [F_(1,24)_ = 174.8; F_(2,48)_ = 4.37; F_(2,48)_ = 89.73; all *p* < 0.001]. Also, sex factor [F_(1,24)_ = 17.69, *p* = 0.0003], time x sex interaction [F_(2,48)_ = 4.37, *p* = 0.018] and treatment x time x sex interaction [F_(2,48)_ = 3.32, *p* = 0.04] were significant (Figure *3A* and B). On day 7 post-injection, MIA-treated mice showed a significant reduction in stride length compared to saline groups in both sexes (*p* < 0.001), with females displaying a stronger impairment than males (*p* < 0.05).By day 28, stride length remained reduced in MIA-treated mice relative to saline groups for both sexes (*p* ≤ 0.01), showing an altered gait, most likely related to functional joint pain.

**Figure 3:**
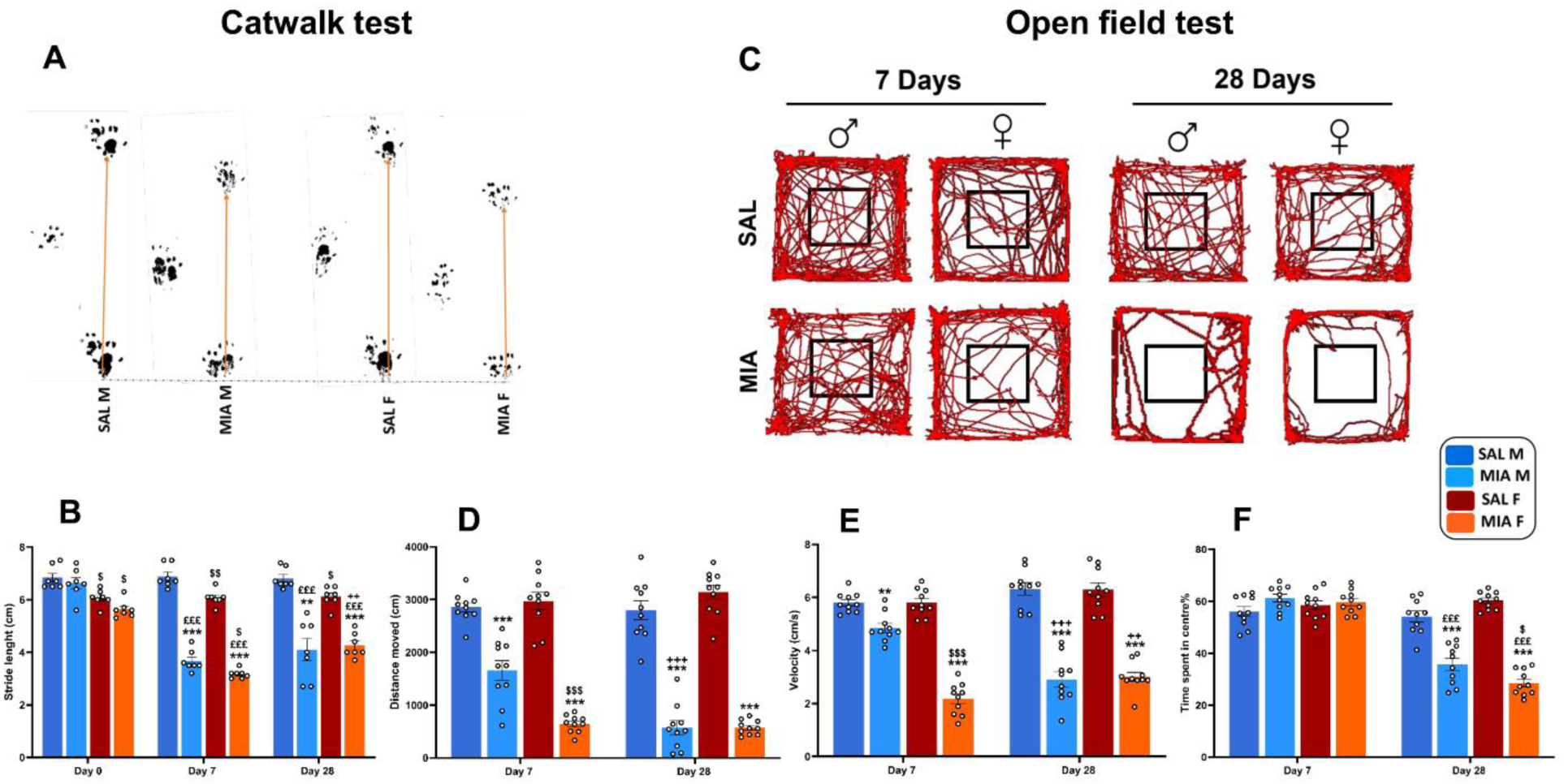
IA injection of MIA induces gait alteration and hypoactivity in a time-dependent way. **A:** Representative images of the stride length decreasing on day 28 post MIA injection. **B:** Stride length. **C:** Representative trajectory tracking of mice in the open field at different time points of the experiment. **D:** Distance moved. **E:** Velocity. **F:** percentage of time spent in the center. Results are represented as mean ± SEM (n=7/group). Holm-Sidak *post hoc*: \*\**p* < 0.01; *** *p* < 0.001 *vs* same sex SAL; £££ *p* < 0.001 *vs* same sex MIA at day 0; ++ *p* < 0.01; +++ *p* < 0.001 *vs* same sex MIA at day 7; $ *p* < 0.05; $$ *p* < 0.01; $$$ *p* < 0.001‘$’ sex difference.

Given the gait alterations observed in osteoarthritic mice, we sought to determine whether these impairments extend to spontaneous locomotor activity. To this end, locomotor behavior in the open field test (Figure *3C, D* and E) was assessed by quantifying total distance traveled (Figure *3D*) and mean locomotor velocity (Figure *3E*).The analysis revealed significant effect of treatment [F_(1, 72)_ = 486.6, *p* < 0.001] and time [F_(1, 72)_ = 7.74, *p* = 0.006] on distance moved. Also, the interactions treatment × time [F_(1, 72)_ = 11.26, *p* = 0.0013], treatment × sex [F_(1, 72)_ = 15.22, *p* = 0.0002], time× sex [F_(1, 72)_ = 10.76, *p* = 0.0016] and treatment × time × sex [F_(1, 72)_ = 4.355, *p* = 0.04] had also a significant effect on distance moved. *Post hoc* analysis using Holm-Sidak’s revealed that MIA IA injection significantly reduced distance moved in MIA-treated males and females compared to their control groups on day 7 (t = 6.35; t = 12.35, *p* < 0.001, respectively). MIA females presenting shorter distance moved compared to males (t = 5.35, *p* < 0.001) consistent with the observed gait alterations. This reductions in mobility persisted on day 28 (t = 11.80; t = 13.61, *p* < 0.001, respectively), with no sex differences associated to this stage (t = 0.01, *p* = 0.98) (Figure *3D*).

Alteration of the mean velocity was also observed (Figure *3E*). Anova analysis showed a significant effect of treatment and sex on animal velocity [F_(1,72)_ = 387.8; F_(1, 72)_ = 20.59, *p* < 0.001]. The interactions treatment × time [F_(1, 72)_ = 13.82, *p* = 0.0004], treatment × sex [F_(1, 72)_ = 19.75, *p* < 0.001], time × sex [F_(1, 72)_ = 23.16, *p* < 0.001] and treatment × time × sex [F_(1, 72)_ = 23.94, *p* < 0.001] were also significant. *Post hoc* analysis showed that MIA IA injection significantly reduced the velocity compared to saline groups in both males (t = 3.31, *p* = 0.004) and females (t = 12.66, *p* < 0.001) on day 7. In addition, MIA females showed lower velocity than MIA males (t = 9.34, *p* < 0.001). The reduction in velocity persisted in both sexes on day 28 (t = 11.93; t = 11.48, *p* < 0.001), but no sex differences were observed at this later time point (t = 0.36, *p* = 0.92) (Figure *3E*).

### 5. IA injection of MIA induces anxio-depressive-like behaviors at a late stage

Beyond motor disturbances, chronic pain is well recognized to impact emotional state and cognitive function. In line with this, analysis of locomotor trajectories in the OF test revealed that MIA-treated mice preferentially explored the periphery while avoiding the central zone, reflecting increased anxiety-like behavior. To determine whether pain-related alterations extended beyond sensory hypersensitivity, additional affective and cognitive domains were examined, including depression-like behavior and working memory. During OF test, MIA-treated mice showed a significant reduction in the percentage of time spent in the center, further indicating enhanced anxiety-like behavior (*Figure 3F*). Three-way ANOVA revealed significant effects of time, treatment, and their interaction on this parameter [F_(1,72)_ = 131.3; F_(1,72)_ = 78.44; F_(1,72)_ = 129.9, *p* < 0.001, respectively], the interaction treatment × sex was also significant [F_(1,72)_ = 11.92, *p* = 0.0009]. *Post hoc* Holm-Sidak’s comparisons revealed no differences in time spent in the center between MIA-treated and control mice on day 7 for either sex (all *p* > 0.05). In contrast, at day 28, MIA mice spent significantly less time in the center compared with controls (t = 7.482; t = 12.77, *p* < 0.001, respectively), with female MIA mice more anxious than males (t = 2.827, *p* = 0.0477). These findings were further confirmed by using the EPM test (*Figure 4A and B*).Three-way ANOVA revealed a significant effect of time, treatment and its interaction on anxiety index [F_(1,72)_ = 31.92; F_(1,72)_ = 17.45; F_(1,72)_ = 29.92, *p* < 0.001, respectively]. Sex factor had also a significant effect [F_(1,72)_ = 7.06, *p* = 0.009 (Figure 4A.B). Consistent with OF results, no differences were observed at day 7, whereas at day 28, MIA-treated male and female mice spent significantly less time in the open arms compared with controls (t = 4.931; t = 6.188, *p* < 0.001, respectively), with females displaying higher anxiety indices than males (t = 2.39, *p* = 0.0380) (*Figure 4A.B*). Together, these results reveal a late onset of anxiety-like behavior in MIA-treated mice, with greater vulnerability in females during the chronic phase of OA pain.

**Figure 4:**
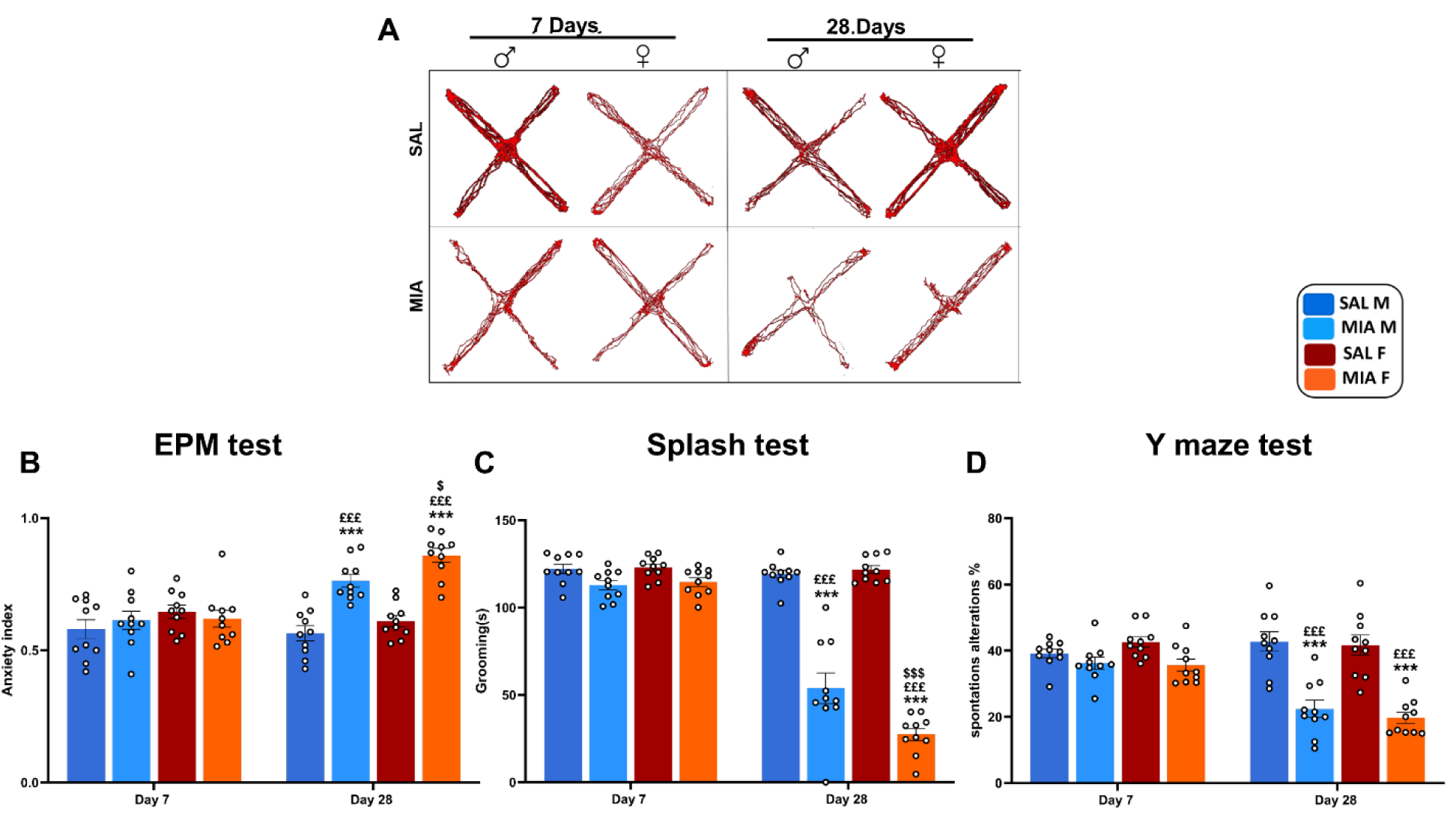
IA injection of MIA induces emotional dysregulation and cognitive impairments in mice. **A:** Representation trajectory tracking of mice EPM test at different time points of the experiment. **B:** Anxiety index. **C:** Grooming time measured during the splash test. **D:** spontaneous alternation was calculated during the Y-maze test. Results are represented as mean ± SEM (n=10/group). Holm-Sidak *post hoc*: *** *p* < 0.001 *vs* same sex SAL; £££ *p* < 0.001 *vs* same sex MIA at day 7; $ *p* < 0.05; $$$ *p* < 0.001 sex difference.

The splash test was then used to measure grooming duration, a behavioral correlate of depression-like behavior in mice. This parameter differed significantly between groups, as revealed by a three-way ANOVA (treatment × sex × time) analysis (Figure *4C*). Results demonstrated a significant effect of both treatment, time, and their interaction on grooming duration [F_(1,72)_ = 256.3; F_(1,72)_ =184.3; F_(1,72)_ = 7.33, *p* < 0.001, respectively]. Also, interactions including sex had a significant effect: treatment × sex [F_(1,72)_ = 6.40, *p* = 0.013]; time × sex [F_(1,72)_ = 5.73, *p* = 0.019]; treatment × time × sex [F _(1,72)_ = 7.33, *p* = 0.008]. Holm-Sidak’s *post hoc* analysis revealed no significant differences in grooming duration between MIA-treated and control mice on day 7 post-injection for either sex (all *p* > 0.05). On day 28, grooming duration decreased in MIA-treated males and females (t = 11.79; t = 17.03, *p* < 0.001), with females exhibiting less grooming duration than males (t= 4.78, *p* < 0.001) (Figure *4C*). These findings demonstrate that depressive-like behaviors emerge at later stages of OA progression and are more pronounced in females.

MIA-treated mice displayed working memory impairments as revealed by the Y maze test (Figure *4D*). Significant effects for treatment, time and its interaction on spontaneous alterations were detected [F_(1,72)_ = 70.81; F_(1,72)_ = 19.05; F_(1,72)_ = 27.33, *p* < 0.001, respectively]. *Post hoc* comparisons showed no significant differences in spontaneous alternations between MIA-treated mice and the control group on day 7 for either sex (all *p* > 0.05). By day 28 post injection, both MIA-treated males and females showed significantly fewer spontaneous alternations compared to their control counterparts (t = 6.55; t = 7.08, *p* < 0.001, respectively). No sex effect was noted (t = 0.88, *p* = 0.61) (Figure *4D*). Overall, these findings demonstrate that working memory impairment emerges only at later stages of OA progression in both sexes.

Behavioral analyses revealed an early onset of joint pain–related behaviors at 7 days post–MIA injection, with a more pronounced effect in female mice during this initial phase of the pathology. These pain-related alterations persisted at 28 days, although sex differences were rebalanced during the chronic stage. Together, these findings validate the MIA-induced osteoarthritis model as a robust model of chronic pain in both sexes. In contrast, anxiety/ depression-like bahavior, and cognitive alterations emerged only at 28 days post-injection and were absent in both sexes during the early phase of the disease.

It is noteworthy that the neuropsychiatric alterations are stronger in females than in males. These observations led us to study the effects of IA injection of MIA on neuronal populations in cerebral areas involved in the sensory–discriminative (AIC) and affective (ACC) components of pain during the early and chronic phases of the disease.

### 6. IA injection of MIA induces decreased neuronal density in ACC and AIC

Neuronal density was reduced in the ACC region in male and female OA mice, at day 28 post MIA injection (Figure *5*A and B). Data analyzed with three-way ANOVA (treatment × sex × time) revealed significant effects of treatment [F _(1,32)_ = 30.35, *p* <0.001], time [F _(1,32)_ = 9.26, *p* = 0.004] and its interaction [F_(1,32)_ = 6.22, *p* = 0.017] on neuronal density. However, sex factor [F_(1,32)_ = 2.45, *p* = 0.12], treatment × sex interaction [F_(1,32)_ = 1.68, *p* = 0.20], time × sex interaction [F_(1,32)_ = 1.64, *p* = 0.20] and treatment × time × sex interaction [F_(1,32)_ = 0.25, *p* = 0.61] were not significant. *Post hoc* comparisons showed no differences among groups on day 7 (all *p* > 0.05). By day 28, MIA injected mice showed reduced neuronal density in ACC versus saline group in males (t = 3.60, *p* = 0.004) and females (t = 4.39, *p* = 0.0006). In MIA mice, ACC contain 14.5 % less neurons in males, and 18.5 % less neurons in females than in control mice, indicating a neuronal loss in ACC during OA progression. No sex differences were noted (t = 1.81, *p* = 0.15) (Figure *5*A and B).

**Figure 5:**
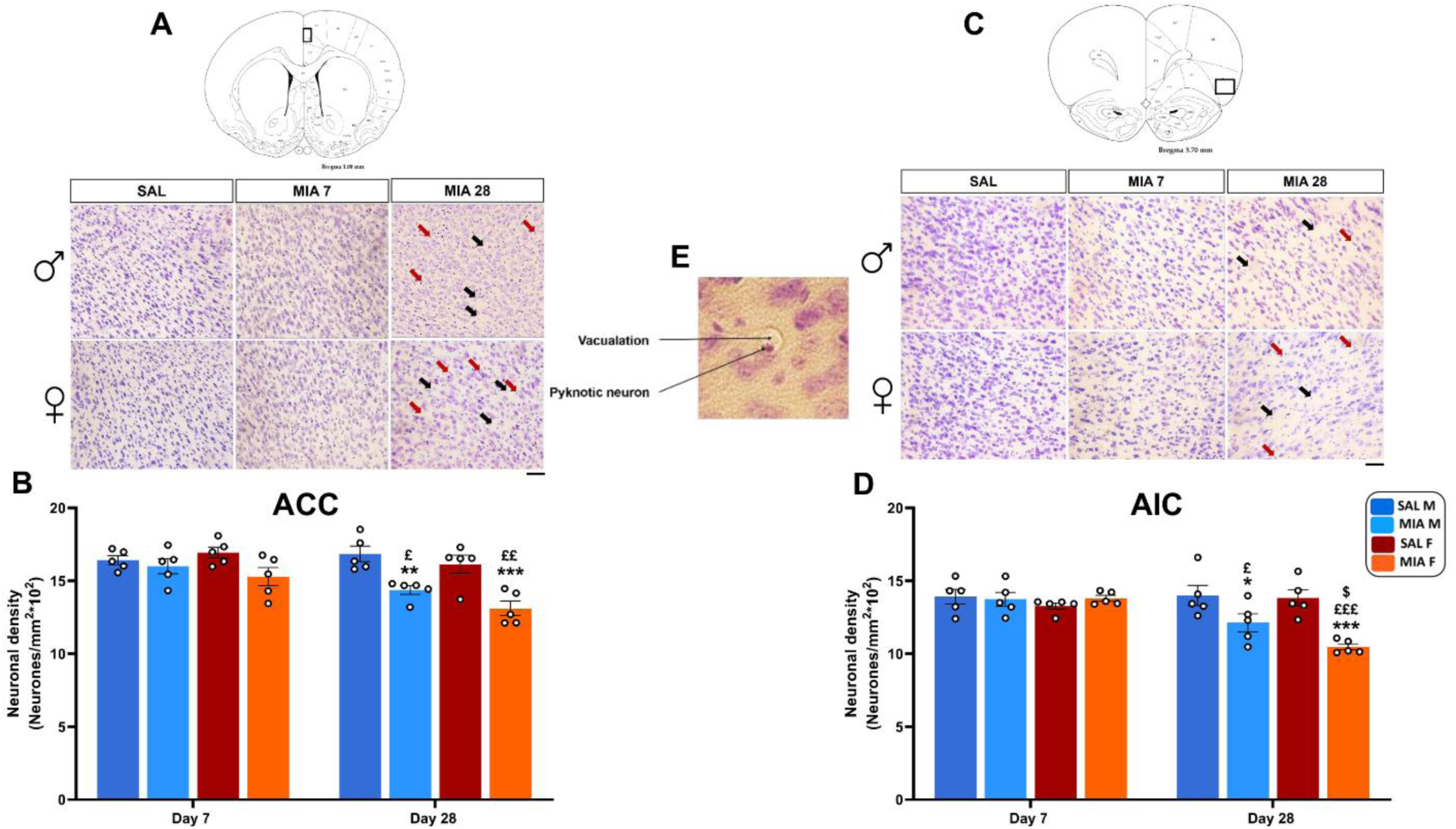
IA injection of MIA leads to neuron loss in ACC and AIC. **A:** Photomicrographs showing the cytoarchitecture of ACC from SAL and MIA male and female mice at different points post MIA injection. **B:** Neuronal density in ACC. **C:** Photomicrographs showing the cytoarchitecture of AIC from SAL and MIA male and female mice at different points post MIA injection. **D:** Neuronal density in AIC. **E:** Schematic, and photomicrograph of pyknotic neuron from ACC at 28 days post-injection. Results are represented as mean ± SEM (n=5/group). Holm-Sidak *post hoc:* \**p* < 0.05; \*\**p* < 0.01; *** *p* < 0.001 *vs* same sex SAL group; £ *p* < 0.05; ££ *p* < 0.01; £££ *p* < 0.001 *vs* same sex MIA at day 7; $ *p* < 0.05 sex difference.

Furthermore, neuronal density was also reduced in the AIC 28 days after MIA treatment (Figure *5*C and D). Three-way ANOVA (treatment × sex × time) showed significant effects of treatment [F_(1,32)_ = 13.43, *p* = 0.0009], time [F_(1,32)_ = 10.51, *p* = 0.002] and their interaction [F_(1,32)_ = 17.43, *p* = 0.0002] on neuronal density. However, sex factor [F_(1,32)_ = 3.29, *p* = 0.07], treatment × sex interaction [F_(1,32)_ = 0.33, *p* = 0.56], time × sex interaction [F_(1,32)_ = 0.90, *p* = 0.34] and treatment × time × sex interaction [F_(1,32)_ = 2.70, *p* = 0.11] were not significant (Figure *5*C and D). *Post hoc* comparisons showed no differences among groups on day 7 (all *p* > 0.05). By day 28, MIA-treated mice showed small but significant reduction with 12.5% of neuronal density in males (t = 2.80, *p* = 0.03) and strong significant reduction of 23.70 % in females (t = 5.03, *p* < 0.001) compared to their control groups. Comparisons withing sex revealed a lower neuronal density in MIA females compared to males (t = 2.49, *p* = 0.04). Although the main effect of sex did not reach significance, *post hoc* comparisons revealed a sex difference at day 28 specifically. Indicating that a large number of AIC neurons are lost in female mice during late OA. Given that the observed degenerating neurons in both regions displayed morphological features consistent with pyknosis, including nuclear condensation, soma shrinkage, and perineuronal vacuolation (Figure *5E*). We further examined apoptotic pathways by analyzing cCasp3^+^ expression and its neuronal co-localization.

### 7. IA injection of MIA induces neuronal loss and increases c-Caspase-3 levels in ACC and AIC

To determine whether chronic OA pain was associated with apoptotic signaling within supraspinal regions, three parameters were analyzed using three-way ANOVA (treatment × time × sex): the density of c-Caspase-3⁻/NeuN⁺ neurons, the density of c-Caspase-3⁺/NeuN⁺ neurons, and the apoptotic index (Figure *6*).

**Figure 6:**
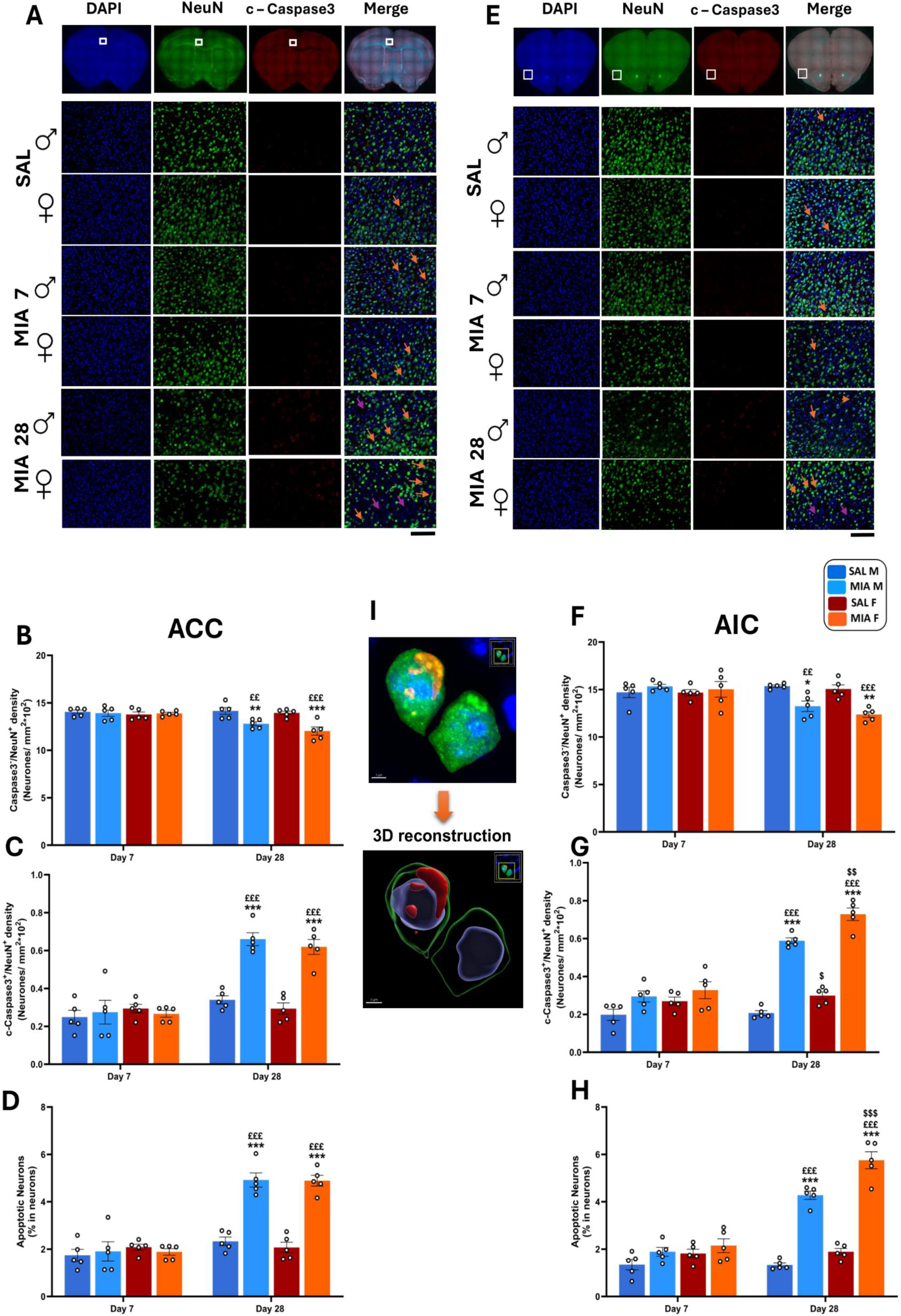
IA injection of MIA induces neuronal loss and increases c-Caspase-3 levels in ACC and AIC. **A:** Photomicrographs showing the cytoarchitecture of ACC from SAL and MIA male and female mice at different points post MIA injection. **B:** c-Caspase-3^-^/NeuN⁺ density in ACC. **C:** c-Caspase-3⁺/NeuN⁺ density in ACC. **D:** Percentage of apoptotic neurons in ACC. **E:** Photomicrographs showing the cytoarchitecture of AIC from SAL and MIA male and female mice at different points post MIA injection. **F:** c-Caspase-3^-^/NeuN⁺ density in AIC. **G:** c-Caspase-3⁺/NeuN⁺ density in AIC. **H:** Percentage of apoptotic neurons in AIC. **I:** 3D reconstruction of an apoptotic neuron from ACC using Imaris showing c-Caspase-3 in cytosol and nucleus (Bleu: DAPI; vert: NeuN; Rouge: c-Caspase-3). Results are represented as mean ± SEM (n=5/group). Holm-Sidak *post hoc*: \**p* < 0.05; ** *p* < 0.01; *** *p* < 0.001 *vs* same sex SAL; ££ *p* < 0.01; £££ *p* < 0.001 *vs* same sex MIA at day 7; $ *p* < 0.05; $$ *p* < 0.01; $$$ *p* < 0.001 sex difference.

In the ACC region the neuronal reduction is confirmed (Figure *6*A and B and C and D), the analysis revealed significant effects of treatment [F_(1,32)_ = 17.57, *p* = 0.0002], time [F_(1,32)_ = 11.76, *p* = 0.0017] and its interaction [F_(1,32)_ = 16.75, *p* = 0.0003] on c-Caspase-3⁻/NeuN⁺ neurons density (Figure *6B*). However, sex factor [F_(1,32)_ = 2.56, *p* = 0.11], treatment × sex interaction [F_(1,32)_ = 0.16, *p* = 0.69], time × sex interaction [F_(1,32)_ = 0.74, *p* = 0.39] and treatment × time × sex interaction [F_(1,32)_ = 1.05, *p* = 0.31] were not significant. *Post hoc* Holm–Sidak comparisons showed no significant differences between groups on day 7 (all *p* > 0.05). By day 28, the analysis revealed significant reductions of c-Caspase-3⁻/NeuN⁺ neurons density in MIA-treated males and females compared to their respective controls (t = 3.42, *p* = 0.006; t = 4.85, *p* = 0.0002, respectively). No sex differences were detected at this stage (t = 1.94, *p* = 0.63) (Figure *6B*). For c-Caspase-3⁺/NeuN⁺ neurons density (Figure *6*C and I), significant effects of treatment, time, and its interaction [F_(1,32)_ = 40.93; F_(1,32)_ = 67.88; F_(1,32)_ =41.53, *p*<0.001] were detected. However, sex factor [F_(1,32)_ = 0.24, *p* = 0.62], treatment × sex interaction [F_(1,32)_ = 0.23, *p* = 0.62], time × sex interaction [F_(1,32)_ = 1.50, *p* = 0.22] and treatment × time × sex interaction [F_(1,32)_ = 0.34, *p* = 0.56] were not significant. *Post hoc* comparisons showed no significant differences between groups on day 7 (all *p* > 0.05). Later, by day 28, significant increase of c-Caspase-3⁺/NeuN⁺ neurons density was noted in both males (65%) and females (58%) MIA-treated mice compared to their control groups (t = 6.37; t = 6.47, *p* < 0.001, respectively). No sex differences were noted at this stage (t = 0.81, *p* = 0.99) (Figure *6*C and I).

Similarly, significant effects of treatment, time and its interaction on apoptotic index were revealed [F_(1,32)_ = 59.05, F_(1,32)_ = 88.85, F_(1,32)_ = 60.48, *p* < 0.001, respectively] (Figure *6D*). *Post hoc* Holm–Sidak comparisons revealed no significant differences between groups at day 7 (all *p* > 0.05). By day 28, MIA-treated mice exhibited a marked increase in the percentage of apoptotic neurons, with levels approximately three times higher in males and four times higher in females compared with their respective control groups (males: t = 7.39; females: t = 8.06, *p* < 0.001). No significant sex difference in the magnitude of apoptosis was observed (t = 0.08, *p* = 0.99; Figure *6D*).

In the AIC (Figure *6*E and F and G and H), the analysis revealed significant effects of treatment [F_(1,32)_ = 8.47, *p* = 0.006] time [F_(1,32)_ = 8.009, *p* = 0.008] and their interaction [F_(1,32)_ = 19.45, *p* = 0.0001] on c-Caspase-3⁻/NeuN⁺ neurons density. However, sex factor [F_(1,32)_ = 1.34, *p* = 0.25], treatment × sex interaction [F_(1,32)_ = 0.42, *p* = 0.52], time × sex interaction [F_(1,32)_ = 0.41, *p* = 0.52] and treatment × time × sex interaction [F_(1,32)_ = 0.04, *p* = 0.82] were not significant (Figure *6F*). *Post hoc* Holm–Sidak comparisons showed no significant differences between groups on day 7 ((all *p* > 0.05). By day 28, the analysis revealed significant reductions of c-Caspase-3⁻/NeuN⁺ neurons density in MIA-treated males and females compared to their respective controls (t = 3.22, *p* = 0.01; t = 4.09, *p* = 0.0013, respectively). No sex differences were detected at this stage (t = 1.33, *p* = 0.34) (Figure *6F*).

c-Caspase-3⁺/NeuN⁺ neurons density increased after MIA injection (Figure *6*G and I). Significant effects of treatment, time, and its interaction [F_(1,32)_ = 154.1, F_(1,32)_ = 88.73, F_(1,32)_ = 71.13, *p* < 0.001] were detected. Also, sex factor significantly affected c-Caspase-3⁺/NeuN⁺ neurons density [F_(1,32)_ = 18.78, *p* = 0.001]. However, treatment × sex interaction [F_(1,32)_ = 0.014, *p* = 0.90], time × sex interaction [F_(1,32)_ = 2.67, *p* = 0.11] and treatment × time × sex interaction [F_(1,32)_ = 1.24, *p* = 0.27] were not significant. *Post hoc* Holm–Sidak comparisons showed no significant differences between groups on day 7 (all *p* > 0.05). By day 28, significant increase of c-Caspase-3⁺/NeuN⁺ neurons density was noted in both males (40 %) and females (51.3 %) MIA-treated mice compared to their control groups (t = 9.80; t = 11.04, *p* < 0.001, respectively). However, MIA-treated females showed higher apoptotic neuronal density compared to MIA-treated males (t = 3.60, *p* = 0.002) (Figure *6*G and I).

Significant effects of treatment, time and sex on apoptotic index were revealed [F_(1,32)_ = 152.5, F_(1,32)_ = 94.91, F_(1,32)_ =19.83, *p* < 0.001, respectively].Also, the interaction treatment × time [F_(1,32)_ = 91.48, *p* < 0.001] and time × sex [F_(1,32)_ = 4.38, *p* = 0.04] were significant (Figure *6H*). *Post hoc* Holm–Sidak comparisons revealed no significant differences between groups at day 7. By day 28, MIA-treated mice exhibited a significant increase in the percentage of apoptotic neurons, with the apoptotic index being approximately twofold higher in males and threefold higher in females compared with their respective control groups (males: t = 9.47; females: t = 12.43, *p* < 0.001).

Notably, MIA-treated females displayed a significantly higher density of c-Caspase-3⁺/NeuN⁺ neurons than MIA-treated males (t = 4.75, *p* < 0.001; Figure *6*H and I). Our results demonstrated sex differences were region-specific, being robust in the AIC but modest or absent in the ACC during late-stage OA.

## Discussion

In this study, we used the MIA-induced OA model in adult male and female mice, a well-established preclinical model widely employed for its ability to replicate the hallmark pathological and symptomatic features of OA (Kuyinu *et al*. 2016; Pitcher *et al*. 2016). Beyond confirming the development of a robust pain phenotype, our work extends previous findings by linking behavioral and affective alterations to region-specific neuronal loss within the ACC and AIC, as evidenced by increased c-Caspase-3 immunoreactivity and reduced neuronal density. Collectively, our results indicate a progressive supraspinal remodeling process, with moderate sex differences, implicating apoptotic mechanisms in the chronic advanced phase of OA pain and its associated comorbidities.

During the early phase of the disease (day 0 to day 7 post–MIA injection), we observed increased mechanical hypersensitivity, reflected by lower mechanical thresholds, together with more pronounced knee swelling in female mice, suggesting a stronger peripheral inflammatory response and enhanced nociceptive sensitivity. These findings are consistent with previous reports demonstrating greater joint inflammation and prolonged hyperalgesia in female rats following MIA injection (Sannajust *et al*. 2019; Ro *et al*. 2020). Such increased vulnerability likely reflects sex-specific neurobiological mechanisms, including estrogen-dependent modulation of nociceptive signaling and immune reactivity (Klein and Flanagan 2019; Chen *et al*. 2021). Despite these early differences, both sexes ultimately exhibited comparable mechanical and thermal hypersensitivity during the advanced stage of OA (day 28), consistent with findings from medial meniscectomy mouse models reporting limited sex effects on evoked pain thresholds during the chronic phase of OA (Temp *et al*. 2020). This supports the interpretation that evoked pain assays primarily reflect peripheral sensitization, whereas sex-related differences may be more pronounced in the affective and cognitive dimensions of chronic pain (Malfait *et al*. 2020).

We acknowledge that these pain measurements, performed at the paw, may not fully reflect the pain phenotype experienced at the affected joint. However, previous studies have shown that joint pathology frequently produces referred pain, whereby nociceptive input originating from the joint is perceived in adjacent cutaneous regions (Mapp *et al*. 2013; Pitcher *et al*. 2016; Krishnan *et al*. 2024). This phenomenon is attributed to peripheral and central sensitization mechanisms that promote the spread of pain hypersensitivity beyond the primary site of injury (Miller *et al*. 2014; Thakur *et al*. 2014). In the present study, pain behavior was assessed on the limb ipsilateral to MIA injection, while neuroanatomical analyses focused on the contralateral ACC and AIC. This experimental approach follows previous studies demonstrating that nociceptive and neuroplastic changes remain predominantly ipsilateral in MIA-induced OA (Miller *et al*. 2014; Thakur *et al*. 2014) and remain stable during the chronic phase at day 28 (Pitcher *et al*. 2016). Our design therefore reflects standard and biologically relevant practice in OA pain research.

Functional impairments, including reduced stride length and locomotor hypoactivity, further validate the pain-related disability induced by MIA. These parameters, which are indicative of spontaneous pain and movement avoidance (Alsalem *et al*. 2020; Çağlar *et al*. 2021), were observed in both sexes but were initially more pronounced in females, reflecting stronger early motivational suppression consistent with our findings of enhanced early inflammation and pain in females, as well as prior MIA studies (Ro *et al*. 2020).

Importantly, sex differences became more evident at later time points in affective comorbidities, as assessed using the elevated plus maze (EPM) and splash tests. By day 28 post–MIA injection, although both sexes exhibited anxiety- and depression-like behaviors, female mice displayed higher anxiety indices and reduced grooming duration. This behavioral pattern mirrors clinical observations reporting a greater prevalence and severity of anxiety and depression in women with chronic musculoskeletal pain (Aaron *et al*. 2025). It also aligns with preclinical evidence demonstrating sex-dependent engagement of limbic circuits, particularly within the ACC and AIC (Liu *et al*. 2020; Haggerty *et al*. 2024). In contrast, Y-maze performance revealed comparable impairments in working-memory function in both male and female MIA-treated mice, indicating that cognitive deficits are similarly affected across sexes.

At the cellular level, our histological analyses provide novel evidence of supraspinal apoptotic signaling in chronic OA pain. At day 28 post–MIA injection, both the ACC and AIC exhibited significant reductions in NeuN-positive neuronal density, increased c-Caspase-3⁺/NeuN⁺ co-labeling, and a higher apoptotic index, with effects being particularly pronounced in the female AIC. These findings are consistent with neuropathic pain models reporting c-Caspase-3–dependent neuronal apoptosis in the spinal dorsal horn (Scholz *et al*. 2005; Inquimbert *et al*. 2018) and in the ACC (Tan *et al*. 2018), and in which caspase inhibition prevents neuronal loss and reduces chronic pain hypersensitivity. Moreover, neuronal remodeling within the ACC and AIC—core hubs of the salience and affective pain network—is likely to exacerbate pain aversion and negative emotional processing in late-stage OA. Rodent studies further demonstrate that interventions restoring cortical synaptic integrity or reducing neuroinflammation can rescue both cortical structure and pain-related behaviors (Miyake *et al*. 2025).

The spatial and temporal pattern of apoptosis observed here parallels human neuroimaging findings reporting gray matter volume reductions in the ACC and insula in patients with chronic OA (Rodriguez-Raecke *et al*. 2009; Alshuft *et al*. 2016; Russell *et al*. 2018). Importantly, some studies report partial reversibility of these alterations following total joint replacement (Rodriguez-Raecke *et al*. 2013). Nevertheless, histological evidence of c-Caspase-3 activation in our study supports the possibility that persistent nociceptive stress may lead to irreversible neuronal loss, particularly in females, whose enhanced pain sensitivity may increase apoptotic vulnerability (Lang and McCullough, 2008).

The increased c-Caspase-3 immunoreactivity observed in cortical neurons is consistent with the dual apoptotic and non-apoptotic roles of caspase-3, including synaptic remodeling and long-term depression (Wang *et al*. 2014). Although c-Caspase-3 localization in NeuN-positive neurons appeared predominantly cytosolic (Figure 6I), additional markers such as cleaved PARP or TUNEL labeling would further substantiate neuronal death (Holota *et al*. 2024). Nevertheless, the concurrent reduction in neuronal density supports apoptosis as a contributor to late-stage cortical remodeling in OA.

Taken together, our results support a two-phase model of OA pain at the supraspinal level, with early peripheral sensitization followed by chronic caspase-dependent neuronal degeneration within affective cortical regions, paralleling spinal apoptotic mechanisms described in neuropathic pain (Scholz *et al*. 2005; Inquimbert *et al*. 2018).

In summary, our study provides novel evidence that chronic OA pain is associated with c-Caspase-3–linked neuronal loss in the ACC and AIC, with sex-dependent vulnerability that is particularly pronounced in the insular cortex of females. Importantly, the inclusion of both males and females in preclinical studies is therefore essential to ensure the translational relevance of experimental findings and to guide the development of sex-informed therapeutic strategies for OA and chronic pain conditions. By bridging human neuroimaging evidence of cortical gray matter alterations with molecular markers of neurodegeneration, these findings highlight the importance of sex-aware, supraspinal approaches for understanding and treating chronic osteoarthritis pain.

## Acknowledgements

We thank Sophie Abelanet for imaging assistance in the Institut de Pharmacologie Moléculaire et Cellulaire (IPMC).

